# Anti Alzheimer’s effects of Neem oil nanoemulsion against Amyloid β_1-40_ induced in human neuroblastoma SH-SY5Y cell line

**DOI:** 10.1101/2024.11.23.625014

**Authors:** Balaji Govindaswamy

## Abstract

Alzheimer’s disease is characterized by cognitive decline associated with neurodegeneration and accumulation of amyloid beta. Here, experimenting with in vitro models of AD using SHSY5Y cell line after exposing them with Aβ_1-40_. The natural derivatives like neem oil, plays a potential role in neuroprotection and reduction of tau aggregates in animal models, eventually becoming a alternative treatment beneficiaries in neurodegenerative disease. Formulating the neem oil into a nanoemulsion with Tween 80® and Brij 30™ for stabilization, can be a beneficial drug development for AD. The optimization of this neem oil nanoemulsion was performed with Response Surface Methodology - Box Behnken Design, showed the S mix 10, Sonication power 55%, and Sonication time 15%; as the optimized value to synthesize a stable neem oil nanoemulsion. Secondly, they subjected to thermodynamic stability analysis for estimating the shelf life duration with different temperature conditions. Amyloid beta induced SHSY5Y tend to show severe cell oxidative stress, to estimate that, the neem oil and neem oil nanoemulsion were subjected to DPPH assay. Further, the neem oil and neem oil nanoemulsion analyzed with the Aβ_1-40_ induced SHSY5Y, resulted in good cytotoxicity, where the nanoemulsion exhibited more reduction in cell viability than neem oil. This trend shows that synthesizing the natural oils into nanocarriers can potentially increase the drug release rate, enhanced stability and good biomedical application

## Introduction

Alzheimer’s disease is a neurological illness that predominantly affects the elderly and is characterised by memory loss, cognitive decline, and behavioural abnormalities. Amyloid β peptide accumulation namely is the pathological hallmark of the illness, resulting in plaque development, neuroinflammation, oxidative stress, mitochondrial dysfunction, and neuronal death [1]. These processes compromise synaptic integrity, which eventually leads to a large loss of neurons. Effective treatment approaches are still elusive despite decades of study, which calls for the development of new medications that might lessen these degenerative aspects of AD [2].

Aβ_1-40_ has a role in metal ion homeostasis, synaptic plasticity, and neuronal repair in physiological settings. However; in pathological circumstances its tendency to misfold and aggregate results in the production of fibrils, soluble oligomers; and eventually amyloid plaques all of which are characteristic elements of AD disease [3,4]. By causing oxidative stress, encouraging neuroinflammation, and compromising synaptic signalling, these aggregated forms degrade cellular processes [5]. The length and aggregation kinetics of Aβ_1-40_ are different from those of its counterpart, Aβ_1-42_. Aβ_1-40_ is the most common type in plasma and cerebrospinal fluid, but Aβ_1-42_ is more hydrophobic and clumps more quickly. According to literature, the course of AD may be significantly influenced by the balance between these two peptides rather than by their individual quantities. Developing Alzheimer’s disease diagnostic indicators and treatment approaches requires an understanding of the pathogenic, structural, and functional characteristics of Aβ_1-40_ [6–8].

*Azadirachta indica* seeds are used to make neem oil, a natural product with a long list of pharmacological benefits that include neuroprotective, anti-inflammatory, and antioxidant actions [9]. Its bioactive components, including gedunin, nimbin, nimbolide, and azadirachtin, have strong anti free radical properties and the capacity to alter important inflammatory pathways [10]. According to literature, these substances may prevent lipid peroxidation in brain tissues, limit microglia activation, and lower the production of proinflammatory cytokines including TNF- and IL-6 of which are linked to the development of AD [11]. Neem oil is a strong contender for anti Alzheimer’s research since it has also been shown in preclinical models to improve mitochondrial function, preserve neuronal survival, and stop Aβ induced cytotoxicity [12]. However, because of its hydrophobic properties, limited bioavailability, and poor cellular penetration, neem oil confronts considerable problems in therapeutic use [13]. Its therapeutic effectiveness is limited by these restrictions, especially when it comes to addressing the central nervous system. Drug delivery methods based on nanotechnology are a workable option by enhancing the pharmacokinetic and pharmacodynamic characteristics of bioactive substances [14–16].

For encapsulating and delivering hydrophobic chemicals, nanoemulsion, a colloidal system made up of oil droplets distributed in water and stabilised by surfactants have become more popular. Hydrophobic chemicals are made more soluble and stable by nanoemulsions, which also shield them from deterioration and facilitate effective cellular absorption. They may penetrate the blood brain barrier, which is a significant challenge in the treatment of CNS illnesses like AD, because of their nano sized droplets [17]. Because encapsulating neem oil into nanoemulsion carriers may efficiently target the central nervous system, they have the potential to be a novel treatment method for AD [18]. Neem oil in its nanoemulsion may lessen oxidative stress, apoptosis and neuroinflammation brought on by Aβ_1-40_, also thought to have anti inflammatory properties in this brain because it can alter pathways including NF, MAPK and JK STAT, which are essential for glial cells to produce proinflammatory mediators. This work is based on optimizing the neem oil nanoemulsion using Response Surface Methodology for a precise nanoscale therapeutic agent that has the capability of suppressing the Aβ_1-40_ induced SH-SY5Y as anti Alzheimer’s model. Furthermore; analyzing them with thermodynamic stability and Antioxidant estimation to check the % of ROS inhibitions and reduction of SH-SY5Y in vitro anti Alzheimer’s activity.

## Materials and methods

### Box Behnken Design using Response Surface Methodology

A response surface methodology in form of Box Behnken design was chosen to statistically optimize the neem oil nanoemulsion for three experimental independent variables Surfactant concentration (w/w %) X1, Sonication power (Watt %) X2, Sonication time (Mins) X3 in three factor levels (-1, 0, +0) table 2; based on the dependent variable effects on particle size (Y1) and Polydispersity index (Y2). Stat ease Design expert software version 23.1 was used to perform the study and for the further statistical analyses. A total of 17 runs in random order were designed by the software to optimize the condition in respect to dependent variables table 1. ANOVA was conducted to confirm the mathematical model, Significance of regression coefficients, coefficient of determination (R2), and the lack of fit was studied for each response. Predicted vs Actual, contour graphs and three-dimensional response surface plots were used to represent the interaction and influence of variables on responses [19,20].

**Table 1:** Experimental design of S mix X1, Sonication power X2 and Sonication time X3.

| Run order | X1 S mix concentration (%) | X2 Sonication power (%) | X3 Sonication time (mins) |
| --- | --- | --- | --- |
| 1 | 15 | 50 | 15 |
| 2 | 10 | 55 | 15 |
| 3 | 10 | 60 | 5 |
| 4 | 10 | 50 | 5 |
| 5 | 10 | 55 | 15 |
| 6 | 15 | 60 | 15 |
| 7 | 10 | 55 | 15 |
| 8 | 10 | 55 | 15 |
| 9 | 15 | 55 | 25 |
| 10 | 5 | 60 | 15 |
| 11 | 10 | 50 | 25 |
| 12 | 5 | 55 | 5 |
| 13 | 5 | 55 | 25 |
| 14 | 10 | 60 | 25 |
| 15 | 15 | 55 | 5 |
| 16 | 5 | 50 | 15 |
| 17 | 10 | 55 | 15 |

**Table 2:** Actual levels at coded factors levels of independent variables.

| Code | Independent variable | Actual levels at coded factor levels |
| --- | --- | --- |

|  |  | -1 | 0 | +1 |
| --- | --- | --- | --- | --- |
| X1 | Surfactant mix (%) | 5 | 10 | 15 |
| X2 | Sonication power (%) | 50 | 55 | 60 |
| X3 | Sonication time (mins) | 5 | 15 | 25 |

### Synthesis of Neem Nanoemulsion

The nanoemulsion was prepared by spontaneous ultrasonication method. Initially, a coarse emulsion formulation was prepared with neem oil (10%), S mix (10%) and aqueous phase (80%) under a magnetic stirrer for 400 rpm and 15 mins at 40°C and was preceded with the ultrasonication. The parameters for the ultra-sonicator (Probe sonicator, Hover labs) were set according to the optimized conditions from Box Behnken design (Run order 5 – S mix – 10% Sonication power 55% and Sonication time 15%). The titanium probe 14 mm diameter was submerged into the emulsions and sonicated according to the conditions set. The synthesized neem nanoemulsion was stored in amber coated bottles at 40°C for further analysis.

### Thermodynamic stability

The optimized neem nanoemulsion was kept in amber coated bottles at different storage conditions i.e. 4°C, 30°C, 45°C and 65°C. The temperature condition studies were noted for 0, 5, 10, 15, 20, and 25 days of storage, the experiment was done in triplicate and measured for particle size and PDI [2].

### Dynamic Light scattering analysis

The particle size was determined by using dynamic light scattering (nanoPartica, Horiba, Japan). The neem nanoemulsion was diluted 100 hold (1:100) to avoid multiple scattering effects, diluted samples were filtered though 2.0mL syringe with 0.2 um syringe filter to avoid bubbles before experiments. For optimization the instrument was stabilized at room temperature (20°C-25°C) for 10 mins [21].

### DPPH scavenging assay

The radical scavenging activity of neem oil and neem nanoemulsion was evaluated using DPPH lipophilic radical scavenging assay. The DPPH solution (0.2 M) was prepared in 95% ethanol, and 1 ml of this solution was added to 2.5ml of neem oil and neem nanoemulsion at different concentrations (0.2 – 1.0 M), and the mixture was kept in dark chamber at ambient temperature for 1 hour. The absorbance was measured at 517 nm using a microplate reader (Biotek, Synergy™, US). The percentage of antioxidant activity was calculated using the following formula [2]:

*Antioxidant activity (Inhibition %) = [(A control - A sample)/A control] x 100*.

### Cell culture

The human neuroblastoma SH-SY5Y cells were maintained in MEM medium supplemented with 10% fetal calf serum, 100 U/ml penicillin and streptomycin in a humid atmosphere of 5% CO_2_ at 38°C. SH-SY5Y cells were plated into 96 well plates. The cells were further exposed with various concentrations of Aβ_1-40_ peptide. The control cells were added with the same medium without Aβ_1-40_.

### Preparation of Amyloid β_1-40_

The Amyloid β_1-40_ was taken in 1.5 mM concentration and incubated in saline solution for 72 hours. After aggregation they were exposed to SH-SY5Y cell lines for further analysis [22,23].

### Cell viability assay using MTT

The protocol was followed with slight modification from [24]; cell viability was assessed using a conventional MTT reduction assay. Amyloid β_1-40_; exposed SH-SY5Y cells at a density of 4 x 10 cells per well placed in 96 well plates with 100 µl of fresh medium supplemented with 10% of FBS. After 24 hour of incubation, the following experimental test design was performed: Group 1: Untreated SH-SHY5Y cells (without Nanoemulsion and Aβ), Group 2: Aβ treated SH-SY5Y cells + 0.2 M of Neem Nanoemulsion, Group 3: Aβ treated SH-SY5Y cells + 0.4 M of Neem Nanoemulsion, Group 4: Aβ treated SH-SY5Y cells + 0.6 M of Neem Nanoemulsion, Group 5: Aβ treated SH-SY5Y cells + 0.8 M of Neem Nanoemulsion, Group 6: Aβ treated SH-SY5Y cells + 1.0 M of Neem Nanoemulsion, Group 7: Aβ treated with SH-SY5Y cells (without nanoemulsion); the same replica was experimented for neem oil from group 2 to group 6 with same concentration. After the treatment period, the culture medium was discarded and 100 µl of MTT (500 µg/ml) was added to all wells and the plates were incubated for 5 hrs. The MTT solution was then removed and 100 µl of DMSO was added to all wells to dissolve the dark blue crystals. The plates were shaken for a few minutes and read on a microplate reader (Biotek, Synergy™, US) using a wavelength of 570 nm. The experiment was done in triplicates [2].

### Statistical analysis

The ANOVA was used to statistically determine the difference in independent variables, along with regression coefficient, contour and 3D response graphs using Stat Ease design expert. The statistical analysis for in vitro assays were performed using Graph Pad Prism software and Origin pro

## Results and Discussion

### Optimisation of Neem oil nanoemulsion using Box Behnken design

Using the probe sonicator, may create a high rise in temperature during the emulsification process. The sonication power and sonication time are important parameters and factors in the process of nanoemulsion synthesis. Increasing the sonication power to a permissible limit can nullify the effects of sediment formation or coalescence, resulting in a transparent and stable nanoemulsion through the experiment. The S mix i.e. Tween 80® and Brij 30™, are widely utilised in food and other products, Tween 80® was used as a primary surfactant in this study due to its low irritation and low toxicity that can minimize droplet size of the nanoemulsions [25,26]. Low limited use Brij 30 will not cause toxicity, and stabilizing the neem nanoemulsion also effectively minimizes the droplet size [27]. ANOVA was used to determine the significance of the coefficients of multiple determinations (R2) and the adjusted coefficient of multiple determination of particle size (Table 2) and PDI (Table 3).

**Table 3:** ANOVA of the regression coefficient of Particle size (Y1)

| ANOVA of Particle size |  | Mean square | F Value | R Value |
| --- | --- | --- | --- | --- |
| Main effect | X1 | 99.90 | 408.72 | < 0.0001 |
|  | X2 | 533.26 | 21566.66 | < 0.0001 |
|  | X3 | 13.86 | 56.01 | < 0.0001 |
| Interaction effects | X1, X2 | 69.56 | 281.09 | < 0.0001 |
|  | X1, X3 | 98.70 | 398.89 | < 0.0001 |
|  | X2, X3 | 904.20 | 3654.10 | < 0.0001 |
| Lack of fit |  | 0.2599 | 1.14 | 0.4722 |
| R<sup>2</sup> |  |  | 0.9986 |  |
| Adjusted R<sup>2</sup> |  |  | 0.9977 |  |
| Predicted R<sup>2</sup> |  |  | 0.9957 |  |

The results from the ANOVA analysis of particle size provide critical insights into how the independent variables influence the particle size 1. Main effects: the surfactant mix has a significant impact (P < 0.0001) as evidenced by (F = 99.90). This can be attributed to the ability of Tween 80® and Brij 30™ to reduce the interfacial tension between the oil and aqueous phases, stabilizing smaller droplets. The surfactant mix ensures the formation of a robust interfacial film around the droplets, preventing coalescence during the high energy sonication [28]. Sonication power had the highest F value 21566.66 indicating its dominant role in particle size reduction. Higher power induces intense acoustic cavitations, generating localized high temperatures and pressures that disrupt large droplets into smaller, more uniform droplets [29]. This process is critical for achieving nanoscale emulsions. Sonication time significantly influenced particle size (P > 0.0001, F=56.01), providing sufficient energy input over time to sustain droplet breakup and stabilization by surfactants. The interaction effects; X2, X3: the strong interaction between sonication power and time (F = 3654.10) highlights their synergistic role in size reduction. Prolonged sonication under high power enables extended cavitations and efficient energy transfer, crucial for achieving smaller and stable droplets [30]. However excessive time can lead to over processing potentially destabilizing the system due to coalescence or micelle formation. X1, X2 the interaction between surfactant mix and sonication power (F = 281.09) underscores the role of surfactants in stabilizing droplets formed under intense cavitations forces. The balance of Tween 80® and Brij 30™ ensures sufficient coverage of the newly formed interfaces during high energy input. X1, X3 this interaction (F = 398.89) suggests that adequate sonication time enhances the surfactants ability to stabilize droplets, as it provides more opportunities for surfactant adsorption at the oil water interface [31].

The PDI a critical measure of size uniformity is significantly influenced by the interplay of X1, X2 and X3 table 4. Surfactant X1: A highly significant linear effect (P < 0.0001, F = 1783.06) demonstrates the importance of the surfactant system in achieving uniform size distribution. Tween 80® and Brij 30™ stabilize the newly formed droplets by forming steric and electrostatic barriers that prevent aggregation [32]. Sonication power X2, as with particle size, sonication power is the most influential variable for PDI, with an F value of 2877.05. Higher power enhances droplet breakup while ensuring that the energy is uniformly distributed throughout the emulsion system. Sonication time X3; prolonged sonication significantly reduced PDI (P < 0.0001, F = 1870.05) by providing adequate energy input for consistent droplet size reduction and stabilization [33]. While the interaction effects of X2X3; the strongest interaction effect (F = 6917.29) reflects the importance of synchronising sonication power and time for uniform size distribution. High power over a controlled time allows for precise droplet size regulation without over processing or introducing instability. X1X2, this interaction (F = 6080.29) reveals the synergistic role of surfactants in stabilizing droplets formed under high power sonication. The efficiency of Tween 80® and Brij 30™ ensures rapid interfacial coverage and stabilization of uniform sized droplets. Overall model performance shows perfect fit values (R^2^, adjusted R^2^, and predicted R^2^ = 1.0000), the quadratic model explains the observed variations in PDI completely. The model underscores the importance of balancing the surfactant composition and sonication conditions to achieve optimal size uniformity. The interplay of the surfactant mix, sonication power, and sonication time highlights their critical roles in stabilizing nanoemulsion with small particle size and uniform distribution [34].

**Table 4:**
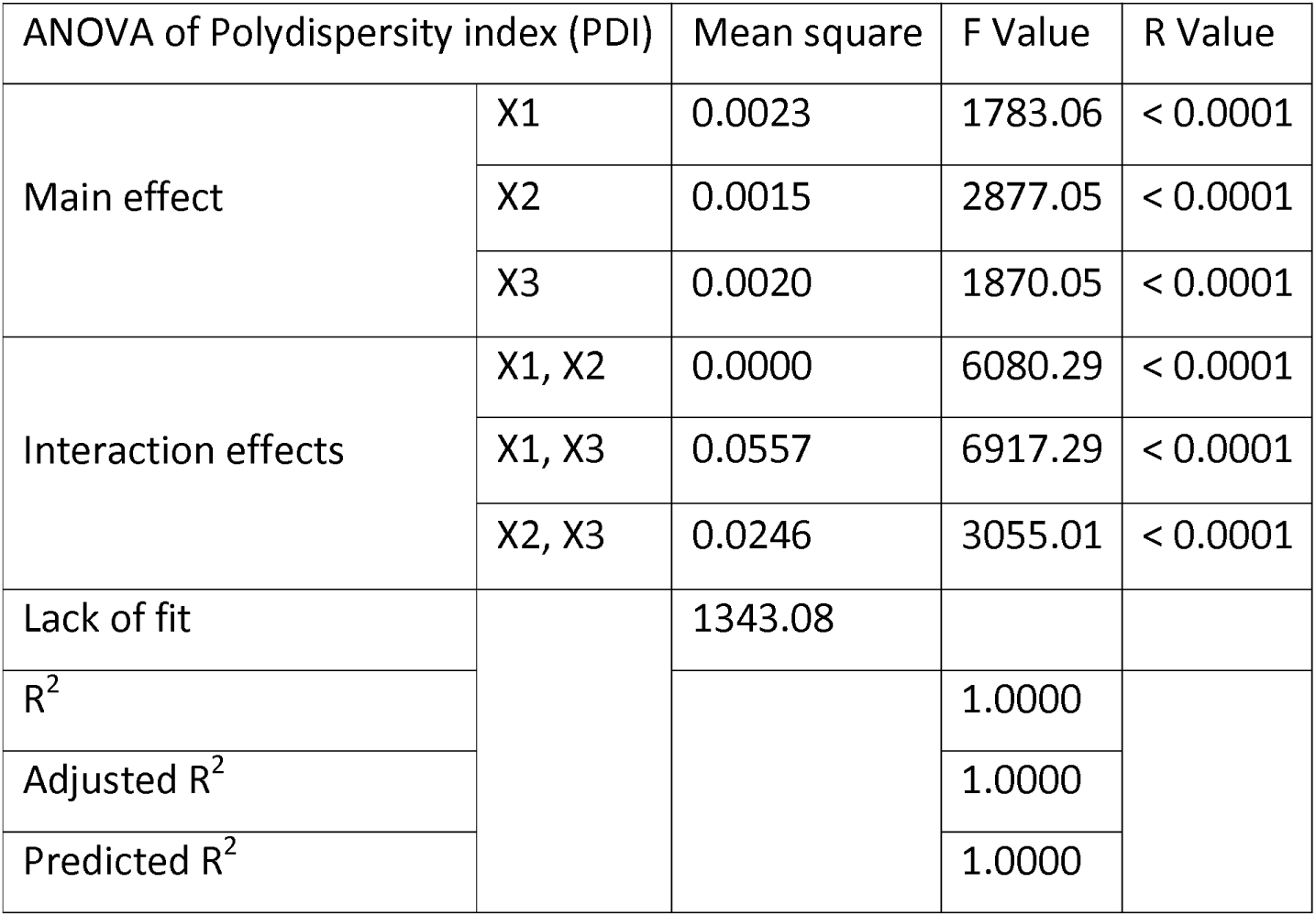
ANOVA of the regression coefficient of Polydispersity index (Y2)

| ANOVA of Polydispersity index (PDI) |  | Mean square | F Value | R Value |
| --- | --- | --- | --- | --- |
| Main effect | X1 | 0.0023 | 1783.06 | < 0.0001 |
|  | X2 | 0.0015 | 2877.05 | < 0.0001 |
|  | X3 | 0.0020 | 1870.05 | < 0.0001 |
| Interaction effects | X1, X2 | 0.0000 | 6080.29 | < 0.0001 |
|  | X1, X3 | 0.0557 | 6917.29 | < 0.0001 |
|  | X2, X3 | 0.0246 | 3055.01 | < 0.0001 |
| Lack of fit |  | 1343.08 |  |  |
| R<sup>2</sup> |  |  | 1.0000 |  |
| Adjusted R<sup>2</sup> |  |  | 1.0000 |  |
| Predicted R<sup>2</sup> |  |  | 1.0000 |  |

The experimental data collected from the responses of the independent variables were analyzed. The response and contour plots of the independent variables on the particle size are shown in fig 1.

**Figure 1:**
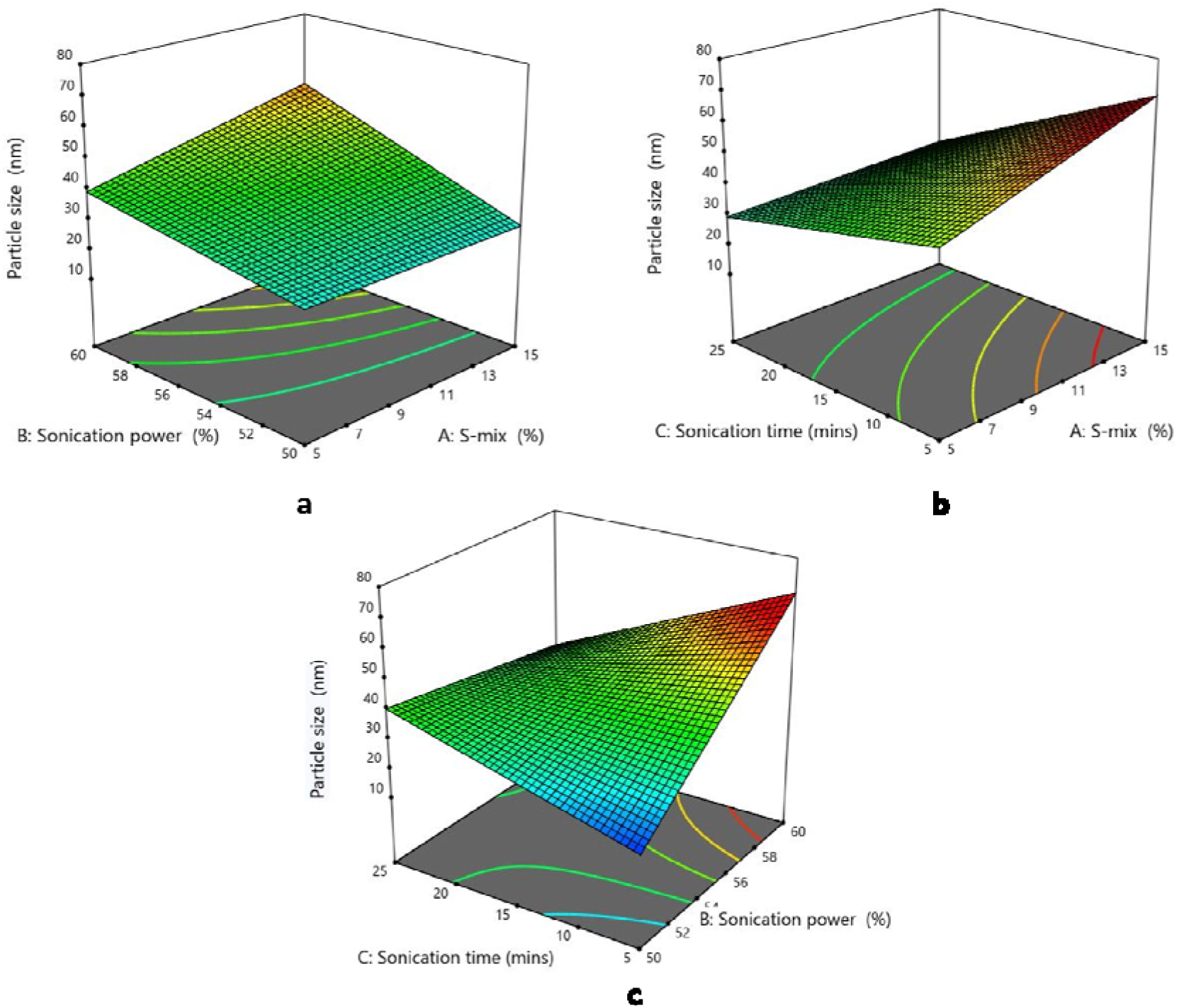
3D response surface interactions of S mix X1, Sonication power X2, and Sonication time X3 in response to Particle size Y1.

Three factor response surface methodology is used to examine how process factors affected particle size. Surfactant mix X1, Sonication power X2, and sonication time X3 were among the variables examined. In Fig 1a; throughout the range of S mix and sonication power investigated, the particle size stays comparatively consistent, according to the response surface plot of particle size as a function of sonication power and S mix [35]. The consistent contour line spacing on the base plane, which shows no interaction between these two elements throughout the studied range, imply that neither sonication power nor S mix significantly affect the particle size on its own in the present experimental set up. Where in Fig 1b; at increasing levels of S mix, the response surface of particle size as a function of sonication time and S mix % shows a larger reliance of particle size on sonication time. Particle size continuously decreases with increasing sonication time; this tendency becomes more noticeable when S mix is raised [36]. According to this pattern, S mix improves stability of smaller particles created by extended sonication, most likely halting re-aggregation. This synergistic impact is supported by the contour lines on the base plane, which show steeper gradients in areas with high S mix, suggesting a more substantial interaction between these variables. In F1c; particle size clearly decreases with increasing sonication time and power, according to the response surface that shows particle size as a function of sonication power and time [37]. The sensitivity of particle size to various factors is shown by the contour lines, which display sharper gradients. The cavitation effect becomes stronger with greater sonication powers, producing enough energy to shatter bigger particles into smaller pieces. Longer sonication time also maintain the cavitation action, enabling more fragmentation. The lowest particle sizes seen are the consequence of this high power and long time combination. These findings highlight how important energy input both in terms of quantity and duration is to producing nanoscale particles [38].

The polydispersity index is critical parameters reflecting the uniformity of particle size distribution, where lower values indicate more monodisperse systems. Fig 2a; the response surface that shows PDI as a function of S mix and sonication power shows very little change in PDI throughout the range of power shows very little change in PDI throughout the range of these two parameters. With consistently spaced contour lines on the base plan and a flat appearance, the surface suggests a modest reliance of PDI on these variables [39]. This indicates that S mix and Sonication power do not significantly affect the systems polydispersity within the studied range. The flat response surface suggests that when these two components are changed separately, PDI stays relatively constant. The intrinsic resilience of the selected formulation in preserving consistent particle size distributions with changing power and stabiliser concentrations may be the cause of this stability [40]. Fig 2b; as the sonication time increase, the PDI significantly drops, especially at greater S mix concentrations, the steep gradient of contour lines at higher S mix levels and the downward sloping surface indicate that the uniformity of particle size distribution is improved by prolonged sonication in conjunction with adequate S mix [41]. Higher S mix concentration stabilise these particles and holds coalescence formation, which leads to more uniform distribution (PDI below 0.500). Longer sonication time; probably supply enough energy to uniformly split particles. This demonstrates how these two parameters majorly impact in maintain the PDI [42]. Fig 2c: in third response surface shows the connection between PDI and the combined effects of sonication power and sonication time. At all sonication power levels, the PDI falls with increasing sonication time; the lowest PDI values are seen at high sonication power and prolonged sonication time [43]. These parameters overwhelming influence is seen in the contour lines, which show a higher gradient along the sonication time axis. The decrease in PDI with increased power and longer sonication time implies that these factors improve particle size homogeneity by encouraging effective cavitation and energy transfer. This leads to better size distribution homogeneity and more consistent particle disintegration [44].

**Figure 2:**
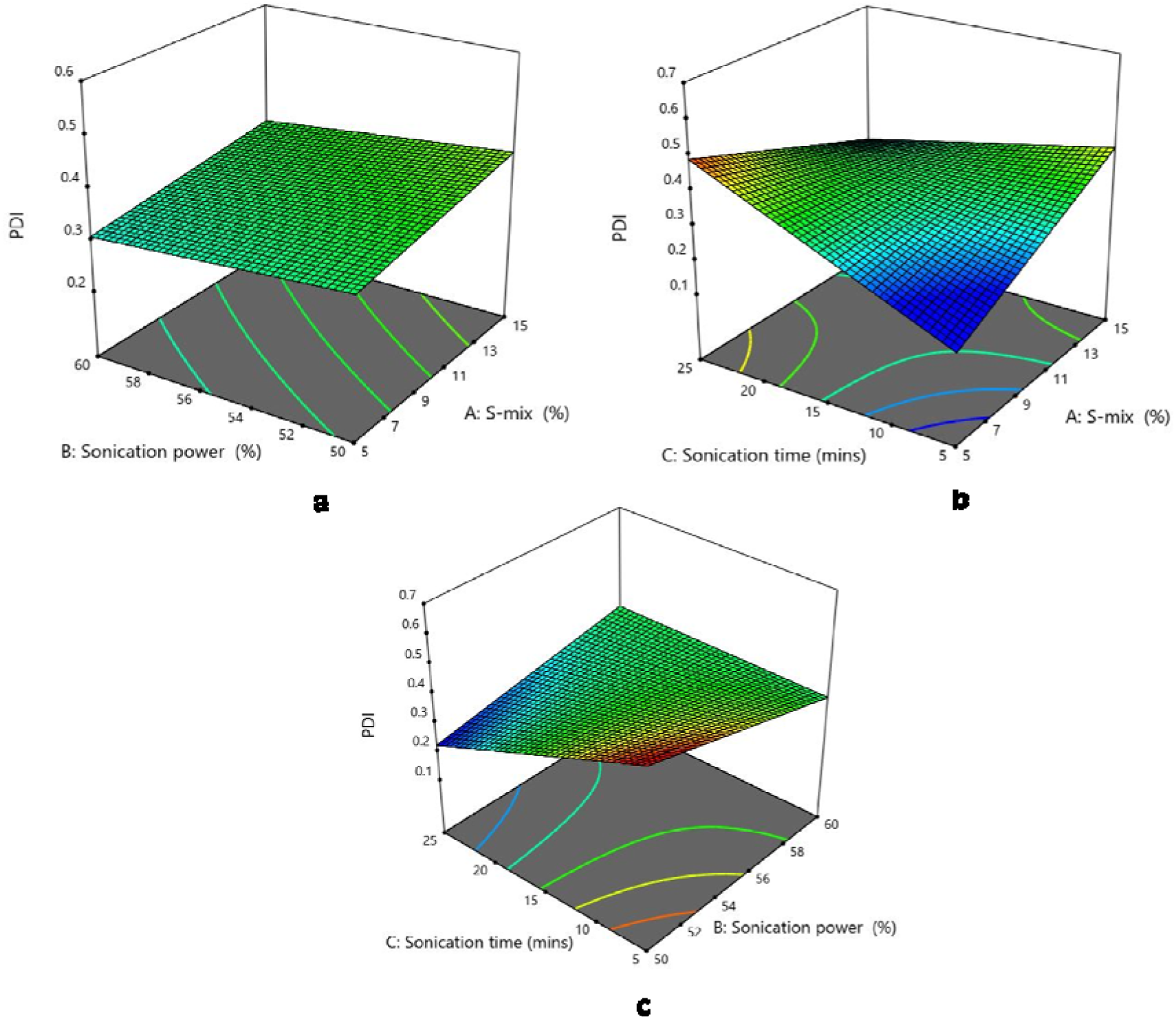
3D response surface interactions of S mix X1, Sonication power X2, and Sonication time X3 in response to PDI Y2.

### Thermodynamic stability and Dynamic Light scattering analysis

The thermodynamic stability of neem oil nanoemulsion was assessed by monitoring changes in particle size over 25 days under storage at four different temperatures (4°C, 30°C, 45°C and 65°C). Fig 3; the results demonstrated a clear influence of temperature on particle size evolution, highlighting the importance of storage conditions for maintaining the stability of the nanoemulsion [45]. At 4°C, the particle size exhibited minimal growth, increasing from 47 nm on the 0^th^ to approximately 63 nm on the 25^th^ day. This indicates that the nanoemulsion retained its structural integrity under refrigerated conditions, likely due to the suppression of thermodynamic process such as molecular diffusion, droplet collision, and coalescence [46]. The stability observed at this temperature underscores the effectiveness of refrigeration in minimizing destabilization phenomena such as Ostwald ripening. When stored at 30°C, the particle size showed a moderate increase from 48 nm on the 0^th^ day to 67 nm on the 25^th^ day. Although the nanoemulsion exhibited acceptable stability at this temperature, the rate of particle size growth was higher than at 4°C. The elevated temperature likely contributed to enhanced droplet mobility and a slight increase in collision frequency, leading to moderate coalescence over time [47]. This suggests that 30°C storage conditions are feasible for short term applications but may compromise long term stability. At 45°C, the particle size increased more significantly, the higher thermal energy at this temperature accelerated destabilization process, such as droplet aggregation and phase separation, resulting in a more growth in particle size [48]. At 65°C, the same growth in particle size was observed which demonstrates that the globule size of nanoemulsion tends to keep expanding as the storage time prolongs [49].

**Fig 3:**
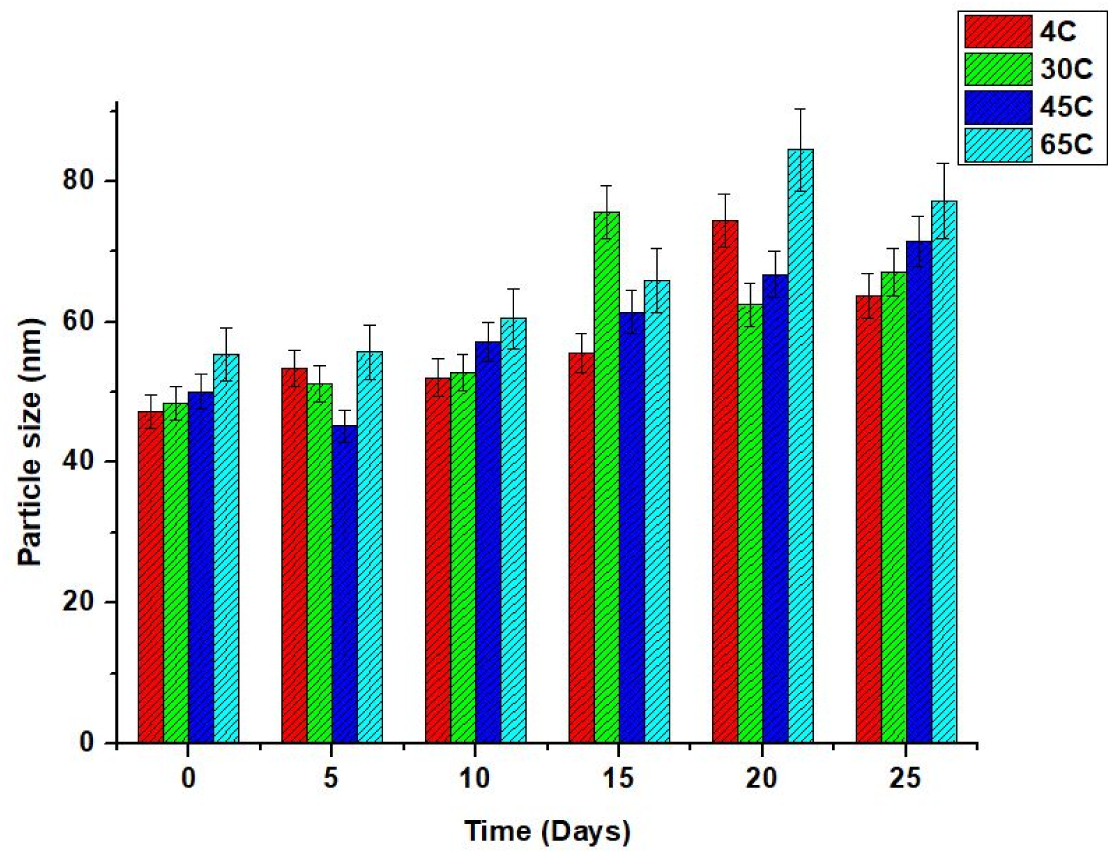
Thermodynamic stability analysis of neem oil nanoemulsion at 4°C, 30°C, 45°C, and 65°C – analyzed for particle size (nm)

The stability of neem oil nanoemulsion was further evaluated by monitoring changes in the polydispersity index (PDI). PDI serves as an indicator of the size uniformity of nanoemulsion droplets, with a value of below 0.5 indicating trends a narrow and uniform size influence [50]. The observed trends in PDI reveal the influence of temperature on the stability of the nanoemulsion. At 4°C, the PDI remained relatively stable throughout the 25^th^ day; this indicates the uniformity of the droplet size distribution by suppressing coalescence [51]. The consistent PDI values at 4°C highlight the high stability of the nanoemulsion under these storage conditions. At 30°C, a similar but slightly higher trend was observed with the PDI increasing from 0.288 to 0.235 on the 25^th^ day, although the nanoemulsions maintained a acceptable level of size uniformity, the higher temperature likely promoted limited coalescence and aggregation of droplets over time, leasing to a gradual increase in heterogeneity [52]. At 45°C, the PDI exhibited a greater degree of fluctuation, increasing from 0.245 to 0.269; the thermal energy enhanced the droplet mobility, increasing the droplet collision and partial coalescence. Meanwhile, at 65°C, the nanoemulsion tends to stay within 0.5 mark line of the PDI standard observatory values. The PDI analysis demonstrates that neem oil nanoemulsion exhibits the highest stability and size uniformity at 4°C. These findings emphasize the importance of proper shelf life conditions in maintaining the physicochemical parameters and stability of neem oil nanoemulsion.

DLS provides valuable information on the homogeneity of the nanoemulsion. Fig 4 shows the DLS graph of optimized neem nanoemulsion, with a particle size of 37 nm. This distribution pattern observed indicates a highly stable and uniform formulation. The nano particle size of the neem oil nanoemulsion is a significant advantage, particularly in applications targeting neurological disorders such as Alzheimer’s disease; it may possess the ability to cross the BBB with the right dosage forms.

**Fig 4:**
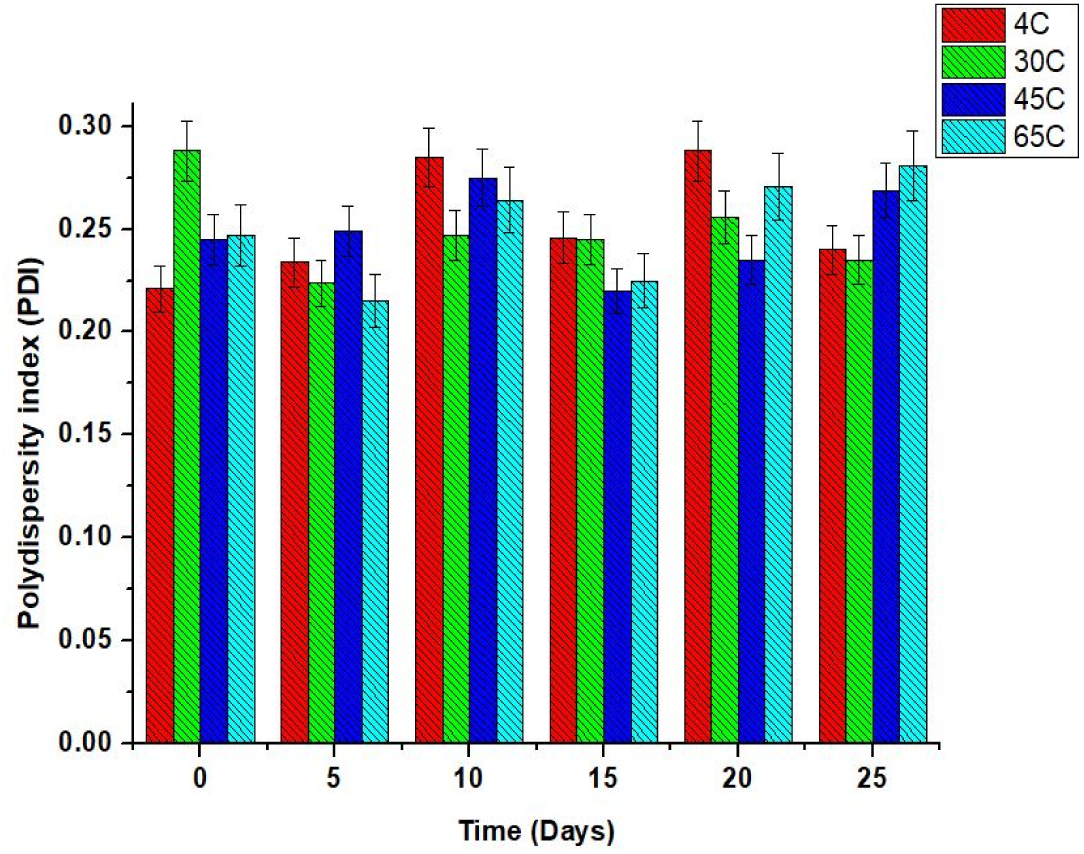
Thermodynamic stability analysis of neem oil nanoemulsion at 4°C, 30°C, 45°C, and 65°C – analyzed for polydispersity index (PDI)

**Fig 5:**
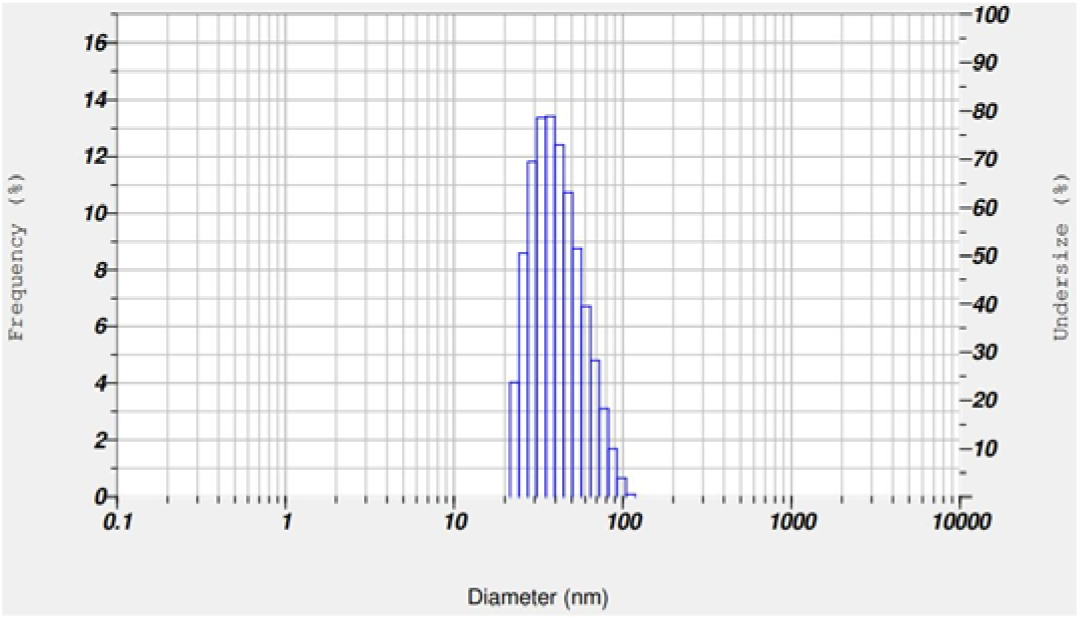
DLS analysis of Neem oil nanoemulsion.

According to Govindaswamy, the nanoemulsions and microemulsions tend to increase the particle size and PDI with more storage durations with different temperatures; this phenomenon will not only cause the emulsion to lose its ability to be intact but might lose its viscous nature, enhancing properties and the efficacy to experiment [2].

**Fig 6:**
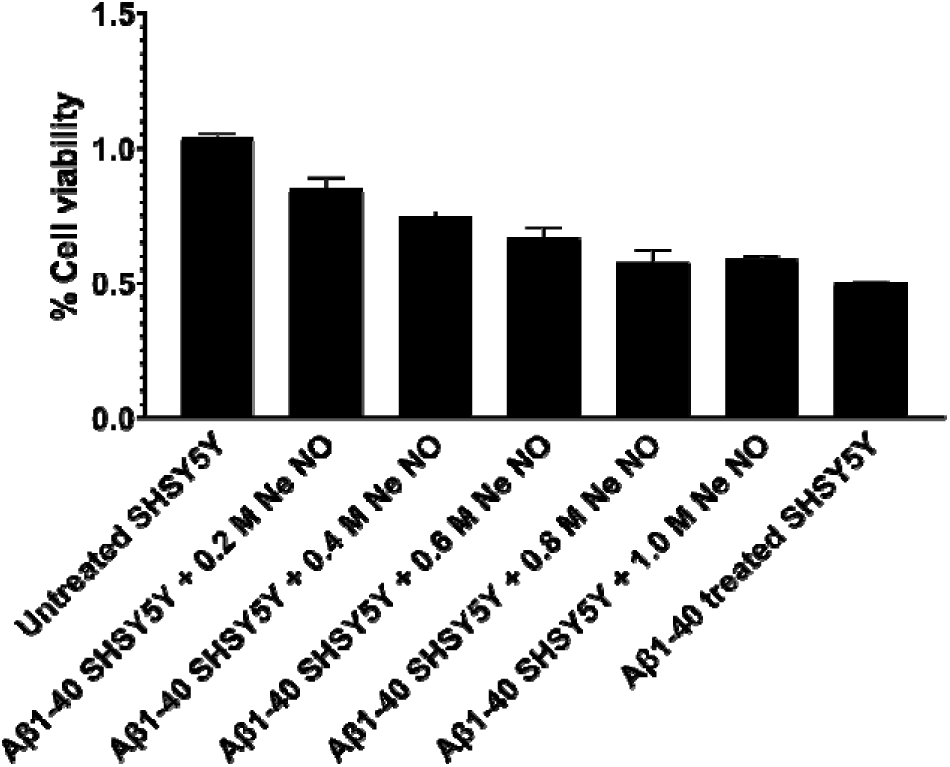
Anti Alzheimer’s activity of Neem oil nanoemulsion in Aβ_1-40_ induced in SH-SY5Y cell lines.

### DPPH antioxidant assay

The investigation of neem oil nanoemulsion was screened for DPPH radical scavenging activity by measuring across a range of concentrations (0.2 – 1.0 M). The results revealed a concentration dependent decrease in absorbance for neem oil and nanoemulsion, indicating their ability to neutralize DPPH free radicals effectively [53]. Table 5, Neem nanoemulsion exhibited high absorbance value compared to the raw oil, suggesting that nano-encapsulating neem oil possesses stronger radical scavenging potential at equivalent concentrations [54]. The higher activity of neem oil nanoemulsion may be attributed to the unique characteristics of nanoemulsion such as enhanced solubility and controlled release properties. While these properties can improve the bioavailability of active components, they may alter the immediate availability of antioxidants in the DPPH assay [55]. The nanoemulsion highlighted its potential as a delivery system for sustained radical scavenging. The observed antioxidant activity has significant implications for neurodegenerative conditions such as Alzheimer’s disease, which is strongly associated with oxidative stress and free radical damage in the brain [56]. Neem oil contains bioactive compounds like azadirachtin and nimbin, known for their antioxidant and anti inflammatory properties, which could counteract the oxidative damage linked to Alzheimer’s disease pathogenesis. Moreover, the nanoemulsion as a carrier offers potential penetration across biological barriers and could enhance the delivery of antioxidant compounds to brain tissues [57]. This highlights the potential as a therapeutic strategy to mitigate the oxidative stress, reduce the symptoms and promote neuronal health in Alzheimer’s disease and other neurodegenerative disorders. According to Sumon Giri, the neem extract showed a good antioxidant capability compared with the standards in different concentrations, suggesting that it can enhance good ROS activity while delivering it in a combination therapy, formulating them into a nano-emulgel can enhance its biomedical activity [58].

**Table 5:** Antioxidant DPPH assay of Neem oil and Neem oil Nanoemulsion.

| Concentration (M) | Neem oil nanoemulsion | Neem oil |
| --- | --- | --- |
| 0.2 | 0.552 ± 0.003 | 0.511 ± 0.006 |
| 0.4 | 0.560 ± 0.004 | 0.497 ± 0.007 |
| 0.6 | 0.534 ± 0.005 | 0.464 ± 0.007 |
| 0.8 | 0.488 ± 0.003 | 0.403 ± 0.006 |
| 1.0 | 0.446 ± 0.004 | 0.371 ± 0.005 |

### In vitro anti Alzheimer’s activity

The cell viability graph illustrates the impact of amyloid beta and neem oil nanoemulsion on SH SY5Y neuroblastoma cells. The untreated SH SY5Y cells, serving as a control, exhibited high viability, indicating the normal physiological state of these cells in culture. In contrast, SH SY5Y treated with amyloid beta showed a significant reduction in cell viability, confirming both cytotoxic and neurotoxic effects of Amyloid beta, which mimics the pathological conditions of Alzheimer’s disease by inducing oxidative stress, mitochondrial dysfunction and apoptosis [59]. When Amyloid beta cells were further exposed to different concentrations of neem oil nanoemulsion, a clear decline in cell viability was observed. Lower concentration of neem oil nanoemulsion caused a moderate reducing in viability, while the higher concentrations showed good cytotoxic activity at these levels. This suggests that the neem oil nanoemulsion compounds interact with the stressed cells in manner that amplifies cell death [60]. The cell viability trend suggest a synergistic effects of neem oil nanoemulsion, in exposing with lower concentrations the cells might counteract the oxidative stress leading to moderate effects. However, when concentration increases the bioactive compounds in neem oil such as nimbolide and nimbin tend to exacerbate mitochondrial dysfunction or disrupt intracellular signalling pathways, further accelerating the apoptosis or necrosis. This aligns with the neurodegeneration mechanism of Amyloid beta toxicity, including calcium dysregulation and pro-inflammatory signalling [61].

**Fig 7:**
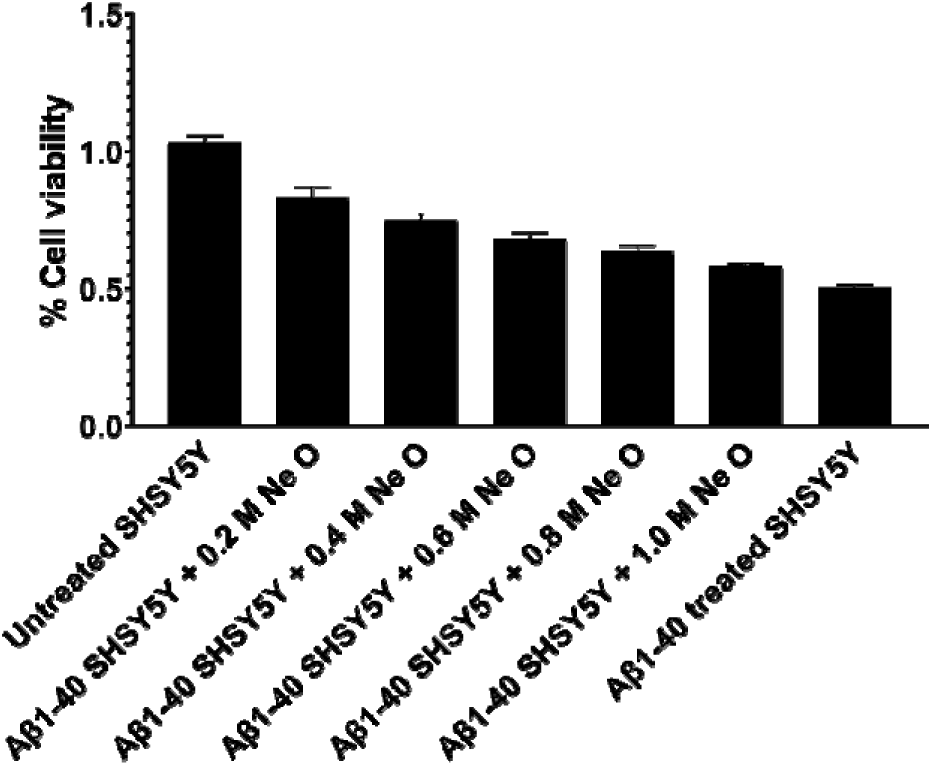
Anti Alzheimer’s activity of Neem oil in Aβ_1-40_ induced in SH-SY5Y cell lines.

The cell viability of neem oil with the same concentrations exhibited similar dose dependent reduction in cell viability of SH SY5Y cells, neem oil 0.2 to 1.0 M showed a progressive cell viability reduction [62]. This reduction indicates that the compounds in neem oil potentiate the cellular stress mechanism activated by amyloid beta. Neem oil is hydrophobic and poorly soluble in aqueous environments, which significantly limits its absorption and systemic bioavailability when consumed or administered directly [63]. Alzheimer’s disease involves the central nervous system, requiring therapeutic agents to cross the Blood Brain Barrier. The raw oil lacks the molecular properties necessary for efficient transport across the BBB. In contrast, nanoemulsion improves the solubility and bioavailability of neem oil by encapsulating its hydrophobic compounds within a nanoscale carrier [64]. These carriers enhance dis-persibility in physiological fluids, allowing better systemic circulation and efficient delivery to the brain. Raw neem oil does not provide a controlled release of its bio actives leading to fluctuations in their plasma concentration. Such variations can reduce therapeutic efficiency or increase the risk of toxicity. Moreover, neem oil active components are prone to degradation in light or gastric fluids, further reducing their effectiveness [65,66].

Nalini states that the study with natural products such as limonoids interacts with the tau protein, leading to degradation and preventing further spreading of neurotoxicity of Amyloid beta. Pointed out the natural compounds like cinnamon, curcumin and nimbin have shown similar effects in preventing tau and amyloid beta aggregation, highlighting the potential of neem derivatives in a broader context of Alzheimer’s treatment [67].

### Conclusion

The optimization experiments for neem nanoemulsion were determined using Box Behnken Design (Response Surface Methodology). The model fitting was performed; the ANOVA and experimental diagnostics were performed to produce optimized neem nanoemulsion. the model suggested run order 5 (S mix – 10% Sonication power 55% and Sonication time 15%) the results suggested that thermodynamic stability of neem nanoemulsion had a good stability over time with different temperatures. the DPPH assay showed a good ROS activity of Neem oil nanoemulsion, showcasing its natural ability to overcome oxidative stress in nanoscale. it is worthy to note that, the neem oil nanoemulsion showed a subsequent cytotoxic activity on Amyloid beta induced SHSY5Y cells, which is dose dependent and enhanced the neem derivatives to neurotoxicity and apoptosis, making the neem oil nanoemulsion as a suitable candidate for the treatment of neurodegenerative diseases.

## Author contribution

Balaji Govindaswamy – Ideation, methodology, investigation, result analysis and draft writing and editing.

## References

[1] Jurcău MC, Andronie-Cioara FL, Jurcău A, Marcu F, Ţiț DM, Pașcalău N, et al. The Link between Oxidative Stress, Mitochondrial Dysfunction and Neuroinflammation in the Pathophysiology of Alzheimer’s Disease: Therapeutic Implications and Future Perspectives. Antioxidants 2022;11:2167. 10.3390/antiox11112167.

[2] Govindaswamy B, Ravikrishnan B, Panda D, Perumal S. In vitro anti cancer, anti inflammatory and anti oxidant activity of Celery oil (Apium graveolens)-Myristic acid based microemulsion system: Characterization and Biomedical application 2024. 10.1101/2024.11.09.622815.

[3] Garza-Lombó C, Posadas Y, Quintanar L, Gonsebatt ME, Franco R. Neurotoxicity Linked to Dysfunctional Metal Ion Homeostasis and Xenobiotic Metal Exposure: Redox Signaling and Oxidative Stress. Antioxid Redox Signal 2018;28:1669–703. 10.1089/ars.2017.7272.

[4] Grasso G, Santoro AM, Lanza V, Sbardella D, Tundo GR, Ciaccio C, et al. The double faced role of copper in Aβ homeostasis: A survey on the interrelationship between metal dyshomeostasis, UPS functioning and autophagy in neurodegeneration. Coord Chem Rev 2017;347:1–22. 10.1016/j.ccr.2017.06.004.

[5] Agostinho P, A. Cunha R, Oliveira C. Neuroinflammation, Oxidative Stress and the Pathogenesis of Alzheimers Disease. Curr Pharm Des 2010;16:2766–78. 10.2174/138161210793176572.

[6] Khan S, Barve KH, Kumar MS. Recent Advancements in Pathogenesis, Diagnostics and Treatment of Alzheimer’s Disease. Curr Neuropharmacol 2020;18:1106–25. 10.2174/1570159X18666200528142429.

[7] Ausó E, Gómez-Vicente V, Esquiva G. Biomarkers for Alzheimer’s Disease Early Diagnosis. J Pers Med 2020;10:114. 10.3390/jpm10030114.

[8] Hampel H, Lista S, Khachaturian ZS. Development of biomarkers to chart all Alzheimer’s disease stages: The royal road to cutting the therapeutic Gordian Knot. Alzheimers Dement 2012;8:312–36. 10.1016/j.jalz.2012.05.2116.

[9] Baby AR, Freire TB, Marques GDA, Rijo P, Lima FV, Carvalho JCMD, et al. Azadirachta indica (Neem) as a Potential Natural Active for Dermocosmetic and Topical Products: A Narrative Review. Cosmetics 2022;9:58. 10.3390/cosmetics9030058.

[10] Chew Y-L, Khor M-A, Xu Z, Lee S-K, Keng J-W, Sang S-H, et al. Cassia alata, Coriandrum sativum, Curcuma longa and Azadirachta indica: Food Ingredients as Complementary and Alternative Therapies for Atopic Dermatitis-A Comprehensive Review. Molecules 2022;27:5475. 10.3390/molecules27175475.

[11] Thakur M, Wang B, Verma ML. Development and applications of nanobiosensors for sustainable agricultural and food industries: Recent developments, challenges and perspectives. Environ Technol Innov 2022;26:102371. 10.1016/j.eti.2022.102371.

[12] Hussain MS, Chaturvedi V, Goyal S, Singh S, Mir RH. An Update on the Application of Nano Phytomedicine as an EmergingTherapeutic Tool for Neurodegenerative Diseases. Curr Bioact Compd 2024;20:e251023222648. 10.2174/0115734072258656231013085318.

[13] Mishra K, Rana R, Tripathi S, Siddiqui S, Yadav PK, Yadav PN, et al. Recent Advancements in Nanocarrier-assisted Brain Delivery of Phytochemicals Against Neurological Diseases. Neurochem Res 2023;48:2936–68. 10.1007/s11064-023-03955-3.

[14] Taly A, Corringer P-J, Guedin D, Lestage P, Changeux J-P. Nicotinic receptors: allosteric transitions and therapeutic targets in the nervous system. Nat Rev Drug Discov 2009;8:733–50. 10.1038/nrd2927.

[15] Upadhyay RK. Drug Delivery Systems, CNS Protection, and the Blood Brain Barrier. BioMed Res Int 2014;2014:1–37. 10.1155/2014/869269.

[16] Abbott NJ. Blood–brain barrier structure and function and the challenges for CNS drug delivery. J Inherit Metab Dis 2013;36:437–49. 10.1007/s10545-013-9608-0.

[17] Marhamati M, Ranjbar G, Rezaie M. Effects of emulsifiers on the physicochemical stability of Oil-in-water Nanoemulsions: A critical review. J Mol Liq 2021;340:117218. 10.1016/j.molliq.2021.117218.

[18] Poovaiah N, Davoudi Z, Peng H, Schlichtmann B, Mallapragada S, Narasimhan B, et al. Treatment of neurodegenerative disorders through the blood–brain barrier using nanocarriers. Nanoscale 2018;10:16962–83. 10.1039/C8NR04073G.

[19] Bhadange YA, Karn A, Saharan VK. Ultrasonically intensified extraction and emulsification of Azadirachtin and its bioactivity analysis for fenugreek crop growth. Chem Eng Process - Process Intensif 2024;199:109748. 10.1016/j.cep.2024.109748.

[20] Barabadi H, Honary S, Ebrahimi P, Alizadeh A, Naghibi F, Saravanan M. Optimization of myco-synthesized silver nanoparticles by response surface methodology employing Box-Behnken design. Inorg Nano-Met Chem 2019;49:33–43. 10.1080/24701556.2019.1583251.

[21] Osman Mohamed Ali E, Shakil NA, Rana VS, Sarkar DJ, Majumder S, Kaushik P, et al. Antifungal activity of nano emulsions of neem and citronella oils against phytopathogenic fungi, Rhizoctonia solani and Sclerotium rolfsii. Ind Crops Prod 2017;108:379–87. 10.1016/j.indcrop.2017.06.061.

[22] Bayes-Genis A, Barallat J, De Antonio M, Domingo M, Zamora E, Vila J, et al. Bloodstream Amyloid-beta (1-40) Peptide, Cognition, and Outcomes in Heart Failure. Rev Esp Cardiol Engl Ed 2017;70:924–32. 10.1016/j.rec.2017.02.021.

[23] Fonseca ACRG, Ferreiro E, Oliveira CR, Cardoso SM, Pereira CF. Activation of the endoplasmic reticulum stress response by the amyloid-beta 1–40 peptide in brain endothelial cells. Biochim Biophys Acta BBA - Mol Basis Dis 2013;1832:2191–203. 10.1016/j.bbadis.2013.08.007.

[24] Amiri M, Braidy N, Aminzadeh M. Protective Effects of Fibroblast Growth Factor 21 Against Amyloid-Beta1–42-Induced Toxicity in SH-SY5Y Cells. Neurotox Res 2018;34:574–83. 10.1007/s12640-018-9914-2.

[25] Cheng M, Zeng G, Huang D, Yang C, Lai C, Zhang C, et al. Advantages and challenges of Tween 80 surfactant-enhanced technologies for the remediation of soils contaminated with hydrophobic organic compounds. Chem Eng J 2017;314:98–113. 10.1016/j.cej.2016.12.135.

[26] Tang J, Wang Y, Wang D, Wang Y, Xu Z, Racette K, et al. Key Structure of Brij for Overcoming Multidrug Resistance in Cancer. Biomacromolecules 2013;14:424–30. 10.1021/bm301661w.

[27] Ferreira SLC, Bruns RE, Ferreira HS, Matos GD, David JM, Brandão GC, et al. Box-Behnken design: An alternative for the optimization of analytical methods. Anal Chim Acta 2007;597:179–86. 10.1016/j.aca.2007.07.011.

[28] Chen W-H, Chiu G-L, Chyuan Ong H, Shiung Lam S, Lim S, Sik Ok Y, et al. Optimization and analysis of syngas production from methane and CO2 via Taguchi approach, response surface methodology (RSM) and analysis of variance (ANOVA). Fuel 2021;296:120642. 10.1016/j.fuel.2021.120642.

[29] Hou T-H, Su C-H, Liu W-L. Parameters optimization of a nano-particle wet milling process using the Taguchi method, response surface method and genetic algorithm. Powder Technol 2007;173:153–62. 10.1016/j.powtec.2006.11.019.

[30] Ataeefard M, Shadman A, Saeb MR, Mohammadi Y. A hybrid mathematical model for controlling particle size, particle size distribution, and color properties of toner particles. Appl Phys A 2016;122:726. 10.1007/s00339-016-0242-1.

[31] Islam Shishir MR, Taip FS, Aziz NAb, Talib RA, Hossain Sarker MdS. Optimization of spray drying parameters for pink guava powder using RSM. Food Sci Biotechnol 2016;25:461–8. 10.1007/s10068-016-0064-0.

[32] Chong W-T, Tan C-P, Cheah Y-K, B. Lajis AF, Habi Mat Dian NL, Kanagaratnam S, et al. Optimization of process parameters in preparation of tocotrienol-rich red palm oil-based nanoemulsion stabilized by Tween80-Span 80 using response surface methodology. PLOS ONE 2018;13:e0202771. 10.1371/journal.pone.0202771.

[33] Raviadaran R, Chandran D, Shin LH, Manickam S. Optimization of palm oil in water nano-emulsion with curcumin using microfluidizer and response surface methodology. LWT 2018;96:58–65. 10.1016/j.lwt.2018.05.022.

[34] Tan SF, Masoumi HRF, Karjiban RA, Stanslas J, Kirby BP, Basri M, et al. Ultrasonic emulsification of parenteral valproic acid-loaded nanoemulsion with response surface methodology and evaluation of its stability. Ultrason Sonochem 2016;29:299–308. 10.1016/j.ultsonch.2015.09.015.

[35] Alzorqi I, Ketabchi MR, Sudheer S, Manickam S. Optimization of ultrasound induced emulsification on the formulation of palm-olein based nanoemulsions for the incorporation of antioxidant β-d-glucan polysaccharides. Ultrason Sonochem 2016;31:71–84. 10.1016/j.ultsonch.2015.12.004.

[36] Koocheki A, Kadkhodaee R. Effect of Alyssum homolocarpum seed gum, Tween 80 and NaCl on droplets characteristics, flow properties and physical stability of ultrasonically prepared corn oil-in-water emulsions. Food Hydrocoll 2011;25:1149–57. 10.1016/j.foodhyd.2010.10.012.

[37] Barber BP, Putterman SJ. Observation of synchronous picosecond sonoluminescence. Nature 1991;352:318–20. 10.1038/352318a0.

[38] Cheong AM, Tan CP, Nyam KL. Emulsifying conditions and processing parameters optimisation of kenaf seed oil-in-water nanoemulsions stabilised by ternary emulsifier mixtures. Food Sci Technol Int 2018;24:404–13. 10.1177/1082013218760882.

[39] Pilong P, Chuesiang P, Mishra DK, Siripatrawan U. Characteristics and antimicrobial activity of microfluidized clove essential oil nanoemulsion optimized using response surface methodology. J Food Process Preserv 2022;46:e16886. 10.1111/jfpp.16886.

[40] Sadeghpour Galooyak S, Dabir B. Three-factor response surface optimization of nano-emulsion formation using a microfluidizer. J Food Sci Technol 2015;52:2558–71. 10.1007/s13197-014-1363-1.

[41] Llinares R, Santos J, Trujillo-Cayado LA, Ramírez P, Muñoz J. Enhancing rosemary oil-in-water microfluidized nanoemulsion properties through formulation optimization by response surface methodology. LWT 2018;97:370–5. 10.1016/j.lwt.2018.07.033.

[42] Musa SH, Basri M, Masoumi HRF, Karjiban RA, Malek EA, Basri H, et al. Formulation optimization of palm kernel oil esters nanoemulsion-loaded with chloramphenicol suitable for meningitis treatment. Colloids Surf B Biointerfaces 2013;112:113–9. 10.1016/j.colsurfb.2013.07.043.

[43] Cunha S, Costa CP, Moreira JN, Sousa Lobo JM, Silva AC. Using the quality by design (QbD) approach to optimize formulations of lipid nanoparticles and nanoemulsions: A review. Nanomedicine Nanotechnol Biol Med 2020;28:102206. 10.1016/j.nano.2020.102206.

[44] Shi Y, Li H, Li J, Zhi D, Zhang X, Liu H, et al. Development, optimization and evaluation of emodin loaded nanoemulsion prepared by ultrasonic emulsification. J Drug Deliv Sci Technol 2015;27:46–55. 10.1016/j.jddst.2015.04.003.

[45] Kumar R, Sinha VR. Preparation and optimization of voriconazole microemulsion for ocular delivery. Colloids Surf B Biointerfaces 2014;117:82–8. 10.1016/j.colsurfb.2014.02.007.

[46] Anjali C, Sharma Y, Mukherjee A, Chandrasekaran N. Neem oil (*Azadirachta indica*) nanoemulsion—a potent larvicidal agent against *Culex quinquefasciatus*. Pest Manag Sci 2012;68:158–63. 10.1002/ps.2233.

[47] Sekar G, Sivakumar A, Mukherjee A, Chandrasekaran N. Probing the interaction of neem oil based nanoemulsion with bovine and human serum albumins using multiple spectroscopic techniques. J Mol Liq 2015;212:283–90. 10.1016/j.molliq.2015.09.022.

[48] Iqbal N, Hazra DK, Purkait A, Agrawal A, Kumar J. Bioengineering of neem nano-formulation with adjuvant for better adhesion over applied surface to give long term insect control. Colloids Surf B Biointerfaces 2022;209:112176. 10.1016/j.colsurfb.2021.112176.

[49] Mossa A-TH, Mohamed RI, Mohafrash SMM. Development of a ‘green’ nanoformulation of neem oil-based nanoemulsion for controlling mosquitoes in the sustainable ecosystem. Biocatal Agric Biotechnol 2022;46:102541. 10.1016/j.bcab.2022.102541.

[50] Sharma R, Kumari A, Singh NS, Singh MK, Dubey S, Iqbal N, et al. Development and stability enhancement of neem oil based microemulsion formulation using botanical synergist. J Mol Liq 2019;296:112012. 10.1016/j.molliq.2019.112012.

[51] Asfour HZ, Alhakamy NA, Alam MS, Al-Rabia MW, Md S. Design of Experiment Navigated Methodical Development of Neem Oil Nanoemulsion Containing Tea Tree Oil for Dual Effect Against Dermal Illness: Ex Vivo Dermatokinetic and In Vivo. J Clust Sci 2023;34:1311–23. 10.1007/s10876-022-02301-x.

[52] Iqbal N, Sharma R, Hazra DK, Dubey S, Kumar N, Agrawal A, et al. Successful utilization of waste cooking oil in Neem oil based fungicide formulation as an economic and eco-friendly green solvent for sustainable waste management. J Clean Prod 2021;288:125631. 10.1016/j.jclepro.2020.125631.

[53] Sharma AD, Chhabra R, Jain P, Kaur I, Chauhan A, Rani R. Preparation, Characterization, and Biological Potential of Nanoemulsion from Rosmarinus officinalis L. Essential Oil. BioNanoScience 2023;13:1955–75. 10.1007/s12668-023-01209-8.

[54] Ghosh V, Sugumar S, Mukherjee A, Chandrasekaran N. Neem (Azadirachta indica) Oils. Essent. Oils Food Preserv. Flavor Saf., Elsevier; 2016, p. 593–9. 10.1016/B978-0-12-416641-7.00067-5.

[55] Sharma AD, Chhabra R, Jain P, Kaur I, Amrita, Bhawna. Nanoemulsions (O/W) prepared from essential oil extracted from *Melaleuca alternifolia*lJ: synthesis, characterization, stability and evaluation of anticancerous, anti-oxidant, anti-inflammatory and antidiabetic activities. J Biomater Sci Polym Ed 2023;34:2438–61. 10.1080/09205063.2023.2253584.

[56] Asgari HT, Es-haghi A, Karimi E. Anti-angiogenic, antibacterial, and antioxidant activities of nanoemulsions synthesized by Cuminum cyminum L. tinctures. J Food Meas Charact 2021;15:3649–59. 10.1007/s11694-021-00947-1.

[57] Revathi T, Thambidurai S. Cytotoxic, antioxidant and antibacterial activities of copper oxide incorporated chitosan-neem seed biocomposites. Int J Biol Macromol 2019;139:867–78. 10.1016/j.ijbiomac.2019.07.214.

[58] Giri S, Chakraborty A, Mandal C, Rajwar TK, Halder J, Irfan Z, et al. Formulation and Evaluation of Turmeric- and Neem-Based Topical Nanoemulgel against Microbial Infection. Gels 2024;10:578. 10.3390/gels10090578.

[59] Coelho BP, Gaelzer MM, Dos Santos Petry F, Hoppe JB, Trindade VMT, Salbego CG, et al. Dual Effect of Doxazosin: Anticancer Activity on SH-SY5Y Neuroblastoma Cells and Neuroprotection on an In Vitro Model of Alzheimer’s Disease. Neuroscience 2019;404:314–25. 10.1016/j.neuroscience.2019.02.005.

[60] Zhang L, Yu H, Zhao X, Lin X, Tan C, Cao G, et al. Neuroprotective effects of salidroside against beta-amyloid-induced oxidative stress in SH-SY5Y human neuroblastoma cells. Neurochem Int 2010;57:547–55. 10.1016/j.neuint.2010.06.021.

[61] De Medeiros LM, De Bastiani MA, Rico EP, Schonhofen P, Pfaffenseller B, Wollenhaupt-Aguiar B, et al. Cholinergic Differentiation of Human Neuroblastoma SH-SY5Y Cell Line and Its Potential Use as an In vitro Model for Alzheimer’s Disease Studies. Mol Neurobiol 2019;56:7355–67. 10.1007/s12035-019-1605-3.

[62] Cesa S, Sisto F, Zengin G, Scaccabarozzi D, Kokolakis AK, Scaltrito MM, et al. Phytochemical analyses and pharmacological screening of Neem oil. South Afr J Bot 2019;120:331–7. 10.1016/j.sajb.2018.10.019.

[63] Amra K, Momin M, Desai N, Khan F. Therapeutic benefits of natural oils along with permeation enhancing activity. Int J Dermatol 2022;61:484–507. 10.1111/ijd.15733.

[64] Bucur MP, Bucur B, Marty J-L, Radu G-L. *In vitro* investigation of anticholinesterase activity of four biochemical pesticides: spinosad, pyrethrum, neem bark extract and veratrine. J Pestic Sci 2014;39:48–52. 10.1584/jpestics.D13-062.

[65] Nikolova G, Ananiev J, Ivanov V, Petkova-Parlapanska K, Georgieva E, Karamalakova Y. The Azadirachta indica (Neem) Seed Oil Reduced Chronic Redox-Homeostasis Imbalance in a Mice Experimental Model on Ochratoxine A-Induced Hepatotoxicity. Antioxidants 2022;11:1678. 10.3390/antiox11091678.

[66] Alahmady NF, Alkhulaifi FM, Abdullah Momenah M, Ali Alharbi A, Allohibi A, Alsubhi NH, et al. Biochemical characterization of chamomile essential oil: Antioxidant, antibacterial, anticancer and neuroprotective activity and potential treatment for Alzheimer’s disease. Saudi J Biol Sci 2024;31:103912. 10.1016/j.sjbs.2023.103912.

[67] Gorantla NV, Das R, Mulani FA, Thulasiram HV, Chinnathambi S. Neem Derivatives Inhibits Tau Aggregation1. J Alzheimers Dis Rep 2019;3:169–78. 10.3233/ADR-190118.

